# Bravo! A Mosquito Antiviral Protein that Restricts Arboviruses through an RRM3-Dependent Nuclear Mechanism

**DOI:** 10.64898/2026.07.16.739008

**Authors:** Elizabeth Walsh, Laura A. St Clair, Claudia Rückert

## Abstract

*Culex quinquefasciatus* mosquitoes are vectors for several medically important viruses, yet their antiviral immune responses remain understudied compared to *Aedes* mosquitoes. The aedine broadly antiviral protein (aBravo) is an RNA-binding protein that was found to inhibit arbovirus replication in *Aedes aegypti*, but the function of the *Cx. quinquefasciatus* Bravo ortholog (CqBravo) remains unknown. Here, we examined the role of CqBravo in antiviral immunity in *Cx. quinquefasciatus*-derived Hsu cells and mosquitoes. We found that CqBravo is also broadly antiviral in *Cx. quinquefasciatus*, reducing replication of La Crosse virus (LACV) and Usutu virus (USUV) in Hsu cells and Sindbis virus (SINV) *in vivo*. CqBravo localized to the nucleus, and truncation and mutational analyses identified an RNA recognition motif (RRM3) critical for nuclear localization and antiviral activity. Quantitative proteomic analysis revealed that full-length CqBravo expression was associated with distinct changes in host protein abundance that were lost following removal of RRM3. Proteins altered by CqBravo expression were enriched for pathways involved in genome maintenance, including DNA repair, DNA metabolism, and ATP-dependent DNA processing. Overall, we demonstrate that CqBravo is an important antiviral factor in *Cx. quinquefasciatus* mosquitoes and may play a broader role in mosquito antiviral immunity across species.

## Introduction

*Culex* species mosquitoes are vectors for several pathogens concerning animal and public health, including West Nile virus (1), Saint Louis encephalitis virus (2), and Usutu virus (USUV) (3). Currently, no vaccines are available for these arboviruses (4). Despite their threat to public health, *Culex* mosquito species remain understudied compared to *Aedes* mosquito species. Understanding the immune processes that influence viral infection within *Culex quinquefasciatus* will be vital for the development of innovative strategies that reduce pathogen spread (5).

Arbovirus infection in mosquitoes often persists throughout adulthood, and many factors are important in controlling virus replication (6, 7). One major antiviral pathway in mosquitoes that helps regulate arbovirus infection is RNA interference (RNAi) (8). RNAi utilizes small RNAs to degrade viral RNA (9). Both the siRNA and piRNA pathways have been identified as antiviral in *Aedes aegypti* (8, 10, 11) and *Cx. quinquefasciatus* (12, 13). However, in *Cx quinquefasciatus* cells, none of the key antiviral proteins (Dicer-2, Argonaute-2, or Piwi4) have been shown to exhibit antiviral activity against any flaviviruses (12–14) and all were specifically not antiviral against Usutu virus (USUV). Despite the importance of antiviral RNAi, the roles of the siRNA and piRNA pathways appear to be virus specific in *Cx. quinquefasciatus*, and our understanding of any truly broadly antiviral proteins remains limited. Recently, the aedine broadly active antiviral protein (aBravo) was identified as antiviral against two alphaviruses, Semliki Forest virus and chikungunya virus, and a flavivirus, Zika virus, in *Ae. aegypti* (15). In Varjak et al. (15), aBravo was initially identified in a Dcr-2 pulldown and was also found to bind Piwi4 and Ago2, but its antiviral activity was independent of functional Dcr-2 (15). While aBravo’s antiviral mechanism remains unclear, the protein has three RNA recognition motifs and one poly-adenylate binding domain (15), indicating that aBravo may broadly be involved in RNA binding activity and may interact with cellular mRNAs. Additionally, in the midgut of *Ae. aegypti* carrying the wAu Wolbachia strain, aBravo was upregulated compared to *Ae. aegypti* not carrying Wolbachia (16). While other mosquito species have orthologs of aBravo, including *Aedes, Culex*, and *Anopheles* species (15), the protein is specific to mosquitoes with no known orthologs in other species.

Here, we focus on the role of *Cx. quinquefasciatus* Bravo (CqBravo) in antiviral responses. We characterized the impact of CqBravo silencing on arbovirus infection in *Cx. quinquefasciatus* cells and mosquitoes. We used an orthobunyavirus (LACV) and a flavivirus (USUV) in *Cx. quinquefasciatus* derived Hsu cells, and an alphavirus (Sindbis virus, SINV) in *Cx.quinquefasciatus* mosquitoes to characterize the impact of CqBravo on arboviruses from separate families. We also examined the importance of individual CqBravo protein domains on antiviral function and cellular localization. Finally, we identified proteomic changes that occur upon CqBravo overexpression.

## Methods

### Cell Lines

*Cx. quinquefasciatus* ovarian-derived Hsu cells (17) were grown in Dulbecco’s Modified Eagle Medium (DMEM; Corning #10-013-CV; Corning, New York, NY, USA) supplemented with 10% FBS and antibiotics (100 units/mL penicillin, 100 μg/mL streptomycin, 5 μg/mL gentamicin), and grown at 27 °C, and 5% CO2.

### Mosquitoes

Mosquito larvae of lab-colonized *Cx. quinquefasciatus* (Orange County, 2016) were kindly provided by Dr. Chris Barker and raised on powdered tetra food. Adult mosquitoes were kept at 26–27 °C with a 16:8 light:dark cycle and 45-55% humidity and were provided with water and sugar *ad libitum*.

### Viruses

The LACV strain R97841d, provided by Brandy Russell at CDC Fort Collins, originated from a human brain in Tennessee in 2012. The USUV strain TMN, provided by Dr. James Weger-Lucarelli (Virginia Tech), was initially isolated in the Netherlands in 2016 from a common blackbird (*Turdus merula*). Virus propagation was conducted on Vero cells, and concentration was achieved through the utilization of the Amicon® Ultra-15 Centrifugal Filter (Sigma-Aldrich, #UFC9010 St. Louis, MO, USA) to attain a high titer stock. Determination of virus stock titers was carried out via standard plaque assay on Vero cells. All stocks used in this study were Vero passage 3 or 4.

### Generation of dsRNA

Gene-specific dsRNA was synthesized using the Megascript RNAi kit (Thermo Fisher Scientific, #AM1626, Waltman, MA, USA) following the manufacturer’s guidelines. In summary, PCR primers containing the T7 promoter sequence were used to amplify CqBravo (Vector Base accession CPIJ003288; NCBI: XM_001844962) from Hsu cDNA, prepared using the High-Capacity cDNA reverse transcription kit (Thermo Fisher Scientific, #4368814, Waltman, MA, USA). As a non-specific control dsRNA, GFP dsRNA was made using a GFP-containing plasmid and T7 primers (Table 1). Target regions underwent PCR amplification utilizing Platinum™ SuperFi II PCR Master Mix (Thermo Fisher Scientific, #12368010 Waltman, MA, USA), with verification conducted via gel electrophoresis. Following purification and concentration using the Zymo DNA-25 clean and concentrator kit (Zymo Research, #D4033; Irvine, CA, USA), dsRNA was generated through *in vitro* transcription using the MEGAScript™ RNAi kit (Thermo Fisher Scientific, AM1626, Waltman, MA, USA). The concentrations and purity of dsRNA were assessed utilizing a Thermo Scientific™ NanoDrop™ One spectrophotometer and gel electrophoresis.

**Table 1.**
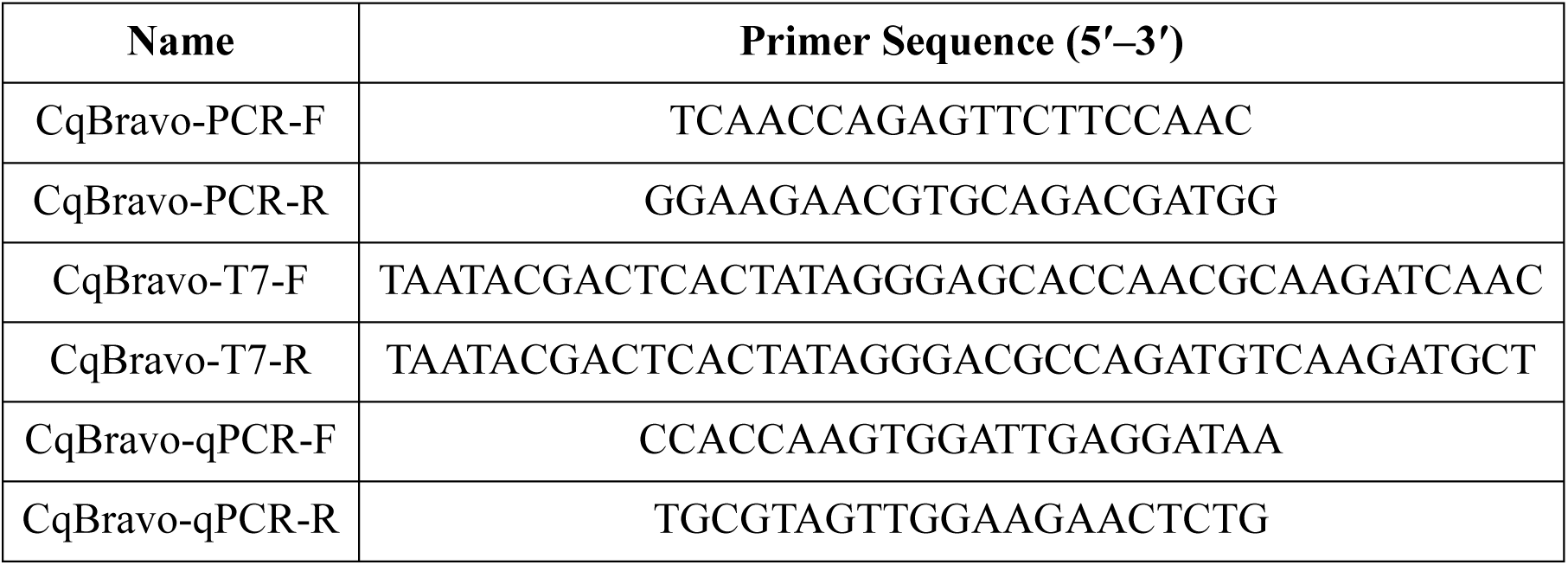
Primers used for CqBravo.

### Transfection of dsRNA

Hsu cells were plated into 24-well plates with 500 µL of culture medium per well. Following overnight incubation at 27°C to ensure cell adhesion, transfection of gene-specific dsRNA, a non-specific dsRNA control (GFP), was carried out using Lipofectamine RNAiMAX (Thermo Fisher Scientific, #13778075, Waltman, MA, USA) as per the manufacturer’s instructions. Briefly, 1.5 µL of Lipofectamine RNAiMAX reagent (Thermo Fisher Scientific, #13778100, Waltman, MA, USA) was diluted in 25 µL Opti-MEM (Thermo Fisher Scientific, #31985062, Waltman, MA, USA) per well. Concurrently, 500 ng of dsRNA was diluted in 25 µL Opti-MEM (Thermo Fisher Scientific, #31985062, Waltman, MA, USA) per well. Subsequently, the diluted dsRNA was added to the diluted Lipofectamine RNAiMAX reagent, mixed thoroughly by pipetting, and incubated for 5 minutes at room temperature. A total of 50 µL of the complex was then added to each well-containing cells, followed by the addition of 450 µL fresh complete media. Cells were further incubated for 48 hours prior to virus infection.

### Virus Infection of Hsu cells

Hsu cells were infected with LACV at an MOI of 10 or USUV at an MOI of 50. These specific MOIs had been previously optimized in our laboratory to infect the most Hsu cells. Briefly, the virus was diluted in DMEM (without any additives) and 250 μL per well was added after the complete culture medium was removed. Following a 2-hour incubation at 27°C to allow for infection, the virus-containing medium was aspirated, and 500 μL of complete culture medium was added to each well. The plates were then incubated at 27°C for 48 hours to measure viral RNA at the peak of replication.

### RNA Extraction

RNA extraction was performed using the Directzol RNA miniprep kit (Zymo Research, #D4033; Irvine, CA, USA) following manufacturer’s instructions. Firstly, for Hsu cells, culture media was aspirated from all wells, and 300 µL of TRI reagent was added to each well. Plates were then placed on a rocker and incubated with TRI reagent for 20 minutes to ensure complete cell lysis. For *Cx. quinquefasciatus* mosquitoes, mosquitoes were cold-anesthetized and placed into homogenization tubes (Benchmark, #D1033-28, Sayreville NJ, USA) with 500 µL of TRI reagent. Mosquitoes were then homogenized using the BeadBug 3 Homogenizer (Benchmark Scientific, #D1030, Sayreville NJ, USA), at a speed of 2800 RPM for 90 seconds. Following incubation or homogenization, cell lysates were transferred to microcentrifuge tubes and stored at -80°C until RNA extraction, following the manufacturer’s protocol. Subsequently, RNA quantification was carried out using a Qubit Flex Fluorometer (Thermo Fisher, Waltham, MA, USA).

### RT-PCR of CqBravo in Mosquito Tissues

To detect whether *CqBravo* mRNA is readily detected, RNA from mosquito life stages and tissues was reverse transcribed into cDNA using the High-Capacity cDNA Reverse Transcription kit (Thermo Fisher Scientific,, #4368814, Waltham, MA USA) and amplified using GoTaq 2x Master Mix (Promega, Madison, WI, USA) and PCR primers listed in Table 1 for 30 cycles of PCR and analyzed on a 1% agarose gel.

### Quantitative Real-Time PCR

Quantitative real-time PCR (qRT-PCR) was performed using either iTaq™ Universal SYBR® Green One-Step Kit (Bio-Rad, #1725150, Hercules, CA, USA) or iTaq Universal Probes 1 step kit (Bio-Rad, #1725141, Hercules, CA, USA) on a CFX96 Touch Real-Time PCR Detection System (Bio-Rad, Hercules, CA, USA). Each target gene or viral RNA was normalized to a previously validated housekeeping gene actin 5 to quantify either gene silencing levels or virus replication as previously described (13). Primers for CqBravo quantification (qPCR) are shown in Table 1.

### Mosquito dsRNA Injections

Concentrated dsRNA was generated similarly to the above description. However, three transcription reactions were performed per gene and pooled to reach a minimum concentration of 5000 ng/µL. Briefly, phenol/chloroform was used to purify and concentrate reactions. For each 50 µL of dsRNA following the RNase/DNase digestion reaction of the Megascript protocol, we added 150 µL of nuclease-free H2O, 100 µL of phenol, and 100 µL of chloroform. The mixture was gently shaken for 15 seconds to avoid foaming and then centrifuged at 15,000 x g for 10 minutes at 4 °C. The top aqueous phase was carefully transferred to a new RNase-free tube, ensuring no interphase or bottom phase transfer. To the aqueous dsRNA, 500 µL of isopropanol was added, followed by incubation at 80 °C for 30 minutes to precipitate RNA. After centrifugation at 15,000 x g for 15 minutes at 4 °C, a visible pellet was observed, and the supernatant was removed. Cold 70% ethanol (600 µL) was added to the pellet, gently pipetted, and then centrifuged at 5,000 x g for 5 minutes at 4 °C. The supernatant was removed, and the pellet was air-dried in a fume hood for 5-10 minutes and then resuspended in a minimal amount of RNase-free water (25-45 µL). A 1:10 dilution of the original product was prepared, and the concentration and purity were assessed using the Nanodrop (Cat#13400525; Thermo Fisher Scientific, Waltham, MA USA).

DsRNA targeting CqBravo, or GFP targeting control dsRNA, was diluted in 1x PBS to a concentration of 5 µg/µl. 0.2 μl of each dsRNA solution was injected into the thorax of cold-anesthetized three-day-old female mosquitoes using a microinjector (Nanoject III, Drummond Scientific Company, PA). Glass capillaries served as injection needles (3.5” Drummond #3-000-203-G/X) (Drummond Scientific Company, PA). The needles were pulled using a Model P-1000 Flaming/Brown Micropipette Puller (Butter Instrument, U.S.) under the following conditions: 430 heat, 50 pull, 75 velocity, and 999 pressure. Following injections, the mosquitoes were placed in cylindrical cardboard containers with a moistened cloth. Mosquitoes were allowed to feed ad libitum on sugar and water following microinjections.

### Mosquito Infectious Bloodmeals

Prior to infection, mosquitoes were starved of sugar for 24 hours and water for 4 hours. Ninety minutes before infection, mosquitoes were placed in a room at 20°C 18-20% humidity to facilitate feeding. Adult female *Cx. quinquefasciatus* mosquitoes 7 days post-eclosion and 3 days post-injection were fed an infectious bloodmeal of defibrinated sheep blood containing 5×10^7^ PFU/mL of SINV isolate 80-2520 and a final concentration of 2 mM ATP. Mosquitoes were allowed to feed for 1 hour on the infected blood meal. Engorged mosquitoes were held for 5 days post-infection (dpi) in a ACL-2 laboratory under the rearing descriptions described above.

### Generation of Bravo Overexpression and Truncation Plasmids

A plasmid encoding GFP under the *Ae. aegypti* polyubiquitin promoter (pUb_eGFP) (13) was used to generate CqBravo constructs and control plasmids. A 3xFLAG-tag sequence from Addgene plasmid #49330 was cloned to the N-terminal of GFP for antibody recognition to create pUb_3xFLAG_GFP. The CqBravo overexpression plasmid (pUb_3xFLAG_CqBravo) was then constructed by replacing the GFP sequence with the CqBravo coding region. The full-length CqBravo coding region was amplified from Hsu cDNA using PCR with Q5® Hot Start High-Fidelity Master Mix (New England Biolabs, #M0515, Ipswich, MA, USA). Truncated CqBravo constructs were generated by amplifying specific regions of the coding sequence. For constructs containing only the C-terminal region of CqBravo, a start codon was added to the truncated sequence to facilitate translation. For the CqBravo-RRM3_mut_, the RNA recognition motif (RRM) was mutated through site-directed mutagenesis. Briefly, oligonucleotide primers were designed with mismatching nucleotides to introduce aromatic ring to alanine mutations into the 3^rd^ RRM (RRM3) region of CqBravo. All plasmid components were amplified via PCR using Q5® Hot Start High-Fidelity Master Mix (New England Biolabs, # M0515, Ipswich, MA, USA).

Subsequently, the PCR products were purified using a Zymoclean Gel DNA Recovery kit (Zymo Research, # D4001, Irvine, CA, USA) and merged into a single plasmid using the NEB Hifi DNA assembly mix (New England Biolabs, #E2621S, Ipswich, MA, USA). The resulting plasmid was introduced into High Efficiency NEB Stable Competent *E. coli* (New England Biolabs, #C3040H, Ipswich, MA, USA) for plasmid amplification. Plasmids were then isolated using the ZymoPURE II Plasmid Maxiprep Kit (Zymo Research, #NC1121573; Irvine, CA, USA). To confirm sequence accuracy, finalized constructs were validated through restriction digestion, Sanger sequencing, and whole plasmid sequencing, performed by Plasmidsaurus using Oxford Nanopore technology with custom analysis and annotation.

### Transfection of Plasmids

Hsu cells were seeded in a 6-well (proteomics), 24-well plate, or optical 96-well plate (microscopy) and allowed to incubate overnight at 27°C and 5% CO_2_. On the day of transfection, 500 μL of fresh media was applied to the cells. The cells were transfected with the plasmid of interest or the control pUb_3xFLAG_EGFP plasmid using X-tremeGENE™ HP DNA transfection reagent (Roche, #6366236001, Burlington, MA, USA) following the manufacturer’s protocol. In brief, the concentration of plasmid DNA was normalized to 100 ng/μL. For each well, X-tremeGENE™ HP DNA transfection reagent was mixed with Plasmid (6-well plate: 2 µg DNA, 24-well plate: 500 ng DNA, 96-well plate: 50 ng) DNA was prepared, using Opti-MEM™ Reduced Serum Medium (Thermo Fisher Scientific, #31985062, Waltham, MA, USA) as a diluent. After a 15-minute incubation period at room temperature, the diluted transfection reagent complex was added to the cells. Mock-transfected cells were treated with X-treme GENE Transfection reagent diluted in Opti-MEM alone. The expression of CqBravo, CqBravo truncations, and CqBravo-RRM3_mut_ was subsequently confirmed through immunostaining using a monoclonal ANTI-FLAG M2 antibody (Sigma Aldrich #F3165, Burlington, MA, USA) at two days post-transfection.

### Immunostaining

For immunostaining, all cells were seeded in 96 well Cellview microplates (Greiner Bio-One, #655891, Monroe, NC). The cells were fixed with 4% paraformaldehyde for 20 minutes, followed by the removal of paraformaldehyde and three washes with 1x PBS for 5 minutes each. Subsequently, the cells were placed in a blocking buffer (1x PBS/5% Normal Goat Serum/0.3% Triton™ X-100) for 1 hour. A primary antibody solution was prepared by diluting the anti-FLAG M2 antibody (Sigma Aldrich #F3165, Burlington, MA, USA) at a ratio of 1:500 in antibody dilution buffer (1x PBS/1% BSA/0.3% Triton™ X-100), which was then added to the cells and incubated overnight at 4 °C. Cells were then washed with 1x PBS three times for 5 minutes each. Following this, a secondary antibody solution containing Alexa Fluor® 555 conjugated anti-mouse IgG (Cell Signaling, #4409S, Danvers, MA, USA) was prepared at a dilution of 1:1000 in antibody dilution buffer and added to the cells. Cells were incubated with secondary antibody for 1 hour at room temperature in the dark, followed by three washes with 1x PBS for 5 minutes each. The cells were counterstained with a 1:10,000 dilution of DAPI and incubated at room temperature for 10 minutes in the dark, followed by a single wash with 1x PBS. Finally, the cells, suspended in 100 µL of 1x PBS, were visualized using a Leica Stellaris 5 HyD S Confocal Microscope.

### Plaque Assays

Vero cells were seeded into 6-well plates and grown to approximately 90% confluency. Virus samples were prepared as ten-fold dilutions in DMEM without fetal bovine serum. After removing the growth medium from each well, 250 µL of the diluted virus was added to the cells and allowed to adsorb for 1.5 hours at 37°C in a 5% CO_2_ incubator. Following adsorption, the inoculum was removed, and 2 mL of a Trag overlay, containing 0.6% Tragacanth gum, 4% FBS, 1x EMEM, 3.5% Sodium Bicarbonate, and 25 mg Gentamicin was added. At the end of the incubation period, the Trag overlay was removed, and cells were stained with 0.1% crystal violet in 20% ethanol for 10–15 minutes. Excess stain was washed off with distilled water, and plates were air-dried. Plaques were visualized and counted to calculate viral titers, expressed as plaque-forming units per milliliter (PFU/mL).

### Proteomics Sample Preparation and Mass Spectrometry

Hsu cells were seeded in 6-well plates and transfected as described above with either pUb_3xFLAG_GFP, pUb_3xFLAG_CqBravo, or pUb_3xFlag_CqBravo_Trunc#2 plasmids. At three days post-transfection, supernatant was removed, cells were washed with 1× PBS, and lysed in 1× RIPA buffer (ThermoFisher, #89900, Waltham, MA, USA) supplemented with Halt™ Protease Inhibitor Cocktail (ThermoFisher, #87785, Waltham, MA, USA) for 20 minutes on ice. Lysates were transferred to microcentrifuge tubes and centrifuged at 13,000 × g for 15 minutes at 4°C. Clarified supernatants were transferred to new tubes, and total protein concentration was determined using the Pierce™ BCA Protein Assay Kit (Thermo Fisher Scientific, #23227, Waltham, MA, USA) according to the manufacturer’s instructions. Samples were normalized to 750 μg/mL in a total volume of 200 μL and stored at −80°C prior to analysis at the University of Nevada, Reno, Mick Hitchcock, Ph.D. Nevada Proteomics Center.

Protein concentrations were re-verified using the fluorescence-based EZQ Protein Quantitation Assay (Invitrogen, #R33200, Carlsbad, CA, USA). Protein extracts were reduced, alkylated with iodoacetamide, and digested using a trypsin/Lys-C protease mixture. Peptides were purified using the EasyPep Mini MS Sample Prep Kit (Thermo Fisher Scientific, #A40006, Waltham, MA, USA) according to the manufacturer’s protocol. Liquid chromatography–mass spectrometry (LC–MS) was performed using an Evosep One LC system (Odense, Denmark) coupled to a Bruker timsTOF Pro 2 mass spectrometer (Billerica, MA, USA) operating in DIA-PASEF mode (18, 19). Peptides were separated using the Evosep 60 samples per day method on a PepSep C18 column (Bruker, #1919826, Billerica, MA, USA). Eluted peptides were ionized using a CaptiveSpray ion source (Bruker, Billerica, MA, USA) and analyzed by mass spectrometry. MS1 spectra were acquired over a mass range of 100–1700 m/z. MS2 acquisition employed a staggered window DIA-PASEF scheme spanning 300–1200 m/z, with 7 PASEF ramps and 25 MS/MS windows across a mobility range of 0.70–1.30 1/K0 and a ramp time of 85 ms, yielding a minimum of seven data points across chromatographic peaks. Raw mass spectrometry data were processed using Spectronaut (Biognosys, Schlieren, Switzerland) in directDIA mode for peptide identification, protein inference, and normalization, using the *C. quinquefasciatus* UniProt reference proteome (UP000002320) and a contaminant database (**c**ommon **R**epository of **A**dventitious **P**roteins, Global Prote**ome** Machine).

### Proteomics Data Processing and Statistical Analysis

Protein-level abundance data generated in Spectronaut (Biognosys, Schlieren, Switzerland) were exported for downstream analysis in R (version 4.5.2) within the RStudio IDE (version 2026.04.0+526, Posit, Boston, MA, USA). Proteins identified as contaminants or decoys and entries lacking quantitative protein abundance values were removed prior to downstream analyses. Protein abundance values were log2-transformed to stabilize variance and approximate normality. Data processing was performed using the dplyr package(20).

Following log2 transformation, PCA and OPLS-DA, and VIP scores were performed using the ropls package(21). VIP plots and corresponding heatmap of protein abundances were generated using ggplot2(22). For differential abundance analysis, mean log2-transformed abundance was calculated per group, and log2 fold change (log2FC) was computed as the difference between group means. Statistical significance was assessed using two-sample t-tests. P-values were adjusted for multiple comparisons using the Benjamini–Hochberg method. Proteins were considered significantly differentially abundant if adjusted p < 0.05 and |log2FC| ≥ 0.5. For statistical comparison at the individual protein level, log2-transformed protein abundances were Z-score normalized within each protein across samples prior to analysis. Z-scores were calculated by centering values to a mean of zero and scaling by the standard deviation within each protein. Z-score values were exported, and violin plots were generated in GraphPad Prism (version 11.0.0, GraphPad Software, Boston, MA, USA). Statistical comparisons for violin plots were performed using one-way ANOVA with Tukey’s post hoc test.

Gene Ontology (GO) enrichment analysis was performed using over-representation analysis with the clusterProfiler package (23). UniProt identifiers were mapped to GO terms using Ensembl Metazoa annotations for *C. quinquefasciatus* (VPISU_Cqui_1.0_pri_paternal). Enrichment was assessed relative to the background of all detected proteins. P-values were adjusted using the Benjamini–Hochberg method, and GO terms with adjusted p < 0.05 were considered significantly enriched.

## Results

### *Cx. quinquefasciatus* Bravo is broadly antiviral against arboviruses *in vitro*

Given that Bravo is antiviral in *Ae. aegypti*, we first wanted to determine the impact of CqBravo on arbovirus infection in *Cx. quinquefasciatus*-derived Hsu cells. Using an RNAi-based method, we silenced CqBravo expression with gene-specific long dsRNA and infected with two arboviruses known to infect Hsu cells, the orthobunyavirus LACV and the flavivirus USUV. Transfection of CqBravo dsRNA resulted in a significant 42% decrease in CqBravo expression (Figure 1A) and LACV was significantly (P <0.001) increased by 2.3-fold when CqBravo was silenced and cells were infected at MOI 1 (Figure 1B). Despite a significant 47% decrease in CqBravo at the time of infection (Figure 1C), there was only a non-significant trend for increased LACV replication following infection at MOI 10 (Figure 1D). For experiments with USUV, we had significant (P<0.05) but modest 37% reduction in CqBravo expression (Figure 1E), which significantly increased (p < 0.05) USUV RNA by 4.8-fold (Figure 1F). Overall, silencing CqBravo expression increased the replication of both arboviruses, suggesting that CqBravo is broadly antiviral in Hsu cells, similar to aBravo activity (15).

**Figure 1.**
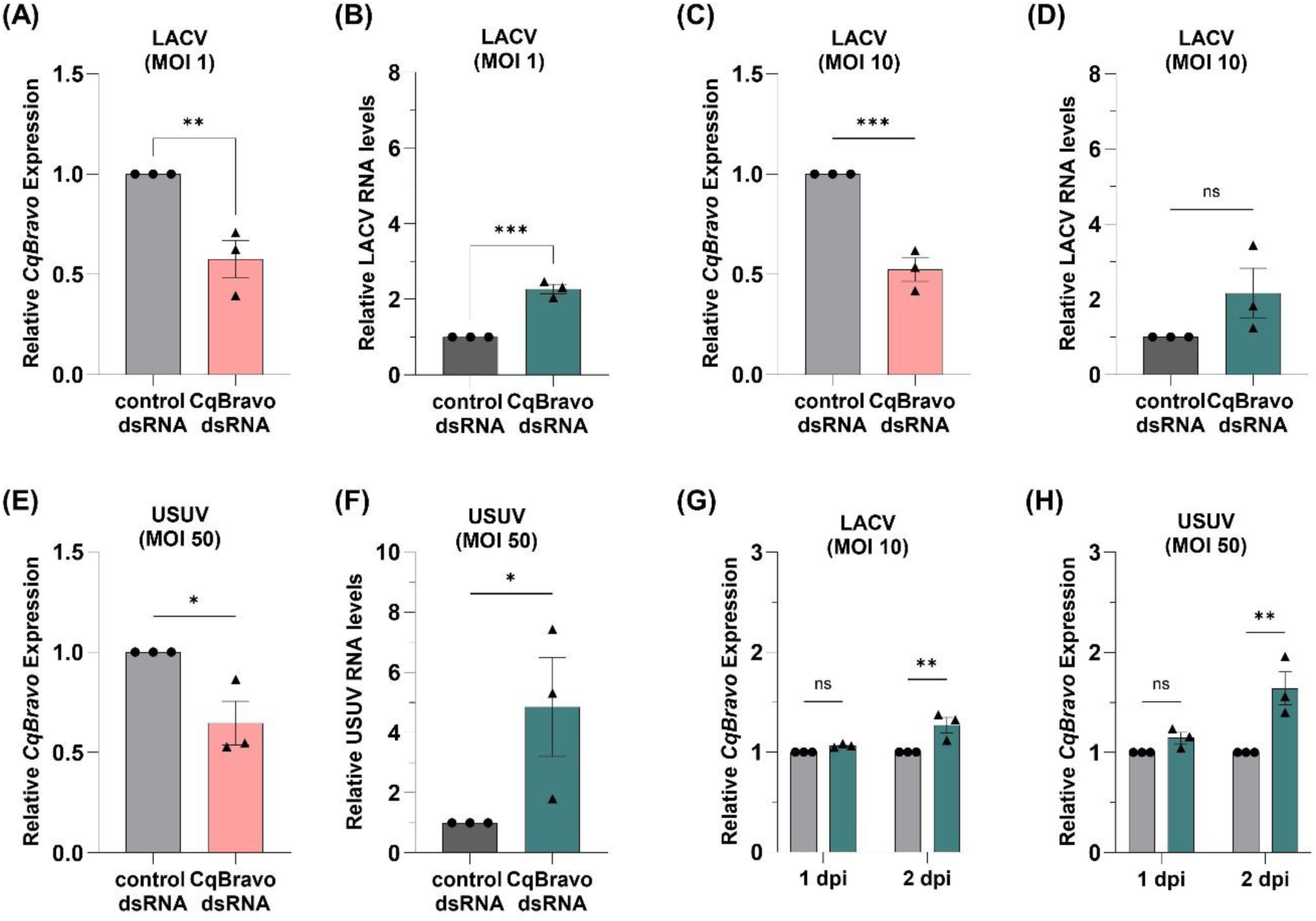
CqBravo is antiviral in *Cx. quinquefasciatus* Hsu cells. Hsu cells were transfected with dsRNA (A-F) targeting CqBravo or GFP as a control. Gene silencing was validated 48 hours post transfection (hpt) by RT-qPCR (A, C, E). Grey bars indicate dsRNA controls (GFP). At 48 hpt, parallel wells of cells transfected with dsRNA were infected with either LACV at MOI 1 (B) or MOI 10 (D) or with USUV at MOI 50 (F). Viral RNA was quantified 2 dpi by RT-qPCR. *CqBravo* expression was also measured in Hsu cells following LACV infection (G) or USUV infection (H) at 1 dpi and 2 dpi. Grey bars indicate uninfected controls, teal bars indicate infected cells (G, H). RNA quantification of *CqBravo*, LACV, and USUV were normalized to *CqActin-5c* and are presented relative to the control GFP dsRNA (A-F) or uninfected control (G, H). Each data point represents one biological replicate experiment with three technical replicates. Bars and error bars indicate the mean of three experiments with SEM. Significant changes in RNA abundance compared to the control were analyzed by one-tailed unpaired t-test (A-F), or two-way ANOVA with Bonferroni correction (G, H). P-values are indicated as *P<0.05, **P<0.01, ***P<0.001, ****P<0.0001.

Next, we determined if arbovirus infection modulates CqBravo expression in Hsu cells. CqBravo expression was measured in Hsu cells infected with either LACV at MOI 10 or USUV at MOI 50 on day 1 or 2 post infection. Both LACV (Figure 1G) and USUV (Figure 1H) infection resulted in a significant increase (P <0.0001, P <0.05, respectively) in CqBravo mRNA by 2 dpi.

However, the very moderate 1.3-fold increase following LACV infection was comparatively less than the 1.6-fold increase following USUV infection. Overall, CqBravo is upregulated in response to arbovirus infection in Hsu cells, indicating a potential inducible antiviral mechanism in *Cx. quinquefasciatus*.

### CqBravo is antiviral in *Cx. quinquefasciatus* mosquitoes

We next looked at *CqBravo* gene expression in *Cx. quinquefasciatus* mosquitoes to determine its role *in vivo*. We first detected *CqBravo* expression during immature life stages of *Cx. quinquefasciatus*, as well as different adult tissues (Figure 2A). We found that *CqBravo* was readily detected in larvae (instar 1-4), pupae, and the head, thorax, abdomen, and midgut of adult females, comparable to the expression of the housekeeping gene *actin* (Figure 2A). We next measured *CqBravo* expression in *Cx. quinquefasciatus* mosquitoes infected via blood meal with the alphavirus SINV. While SINV does not replicate well in Hsu cells, *Culex* mosquitoes are a key vector for it and allow us to extend our research to another group of arbovirus (24). At 3 dpi, there was a trend but no significant increase in *CqBravo* expression, but by 5 dpi *CqBravo* expression was significantly increased (P <0.0001) by 3-fold in mosquitoes infected with SINV compared to uninfected mosquitoes (Figure 2B).

**Figure 2.**
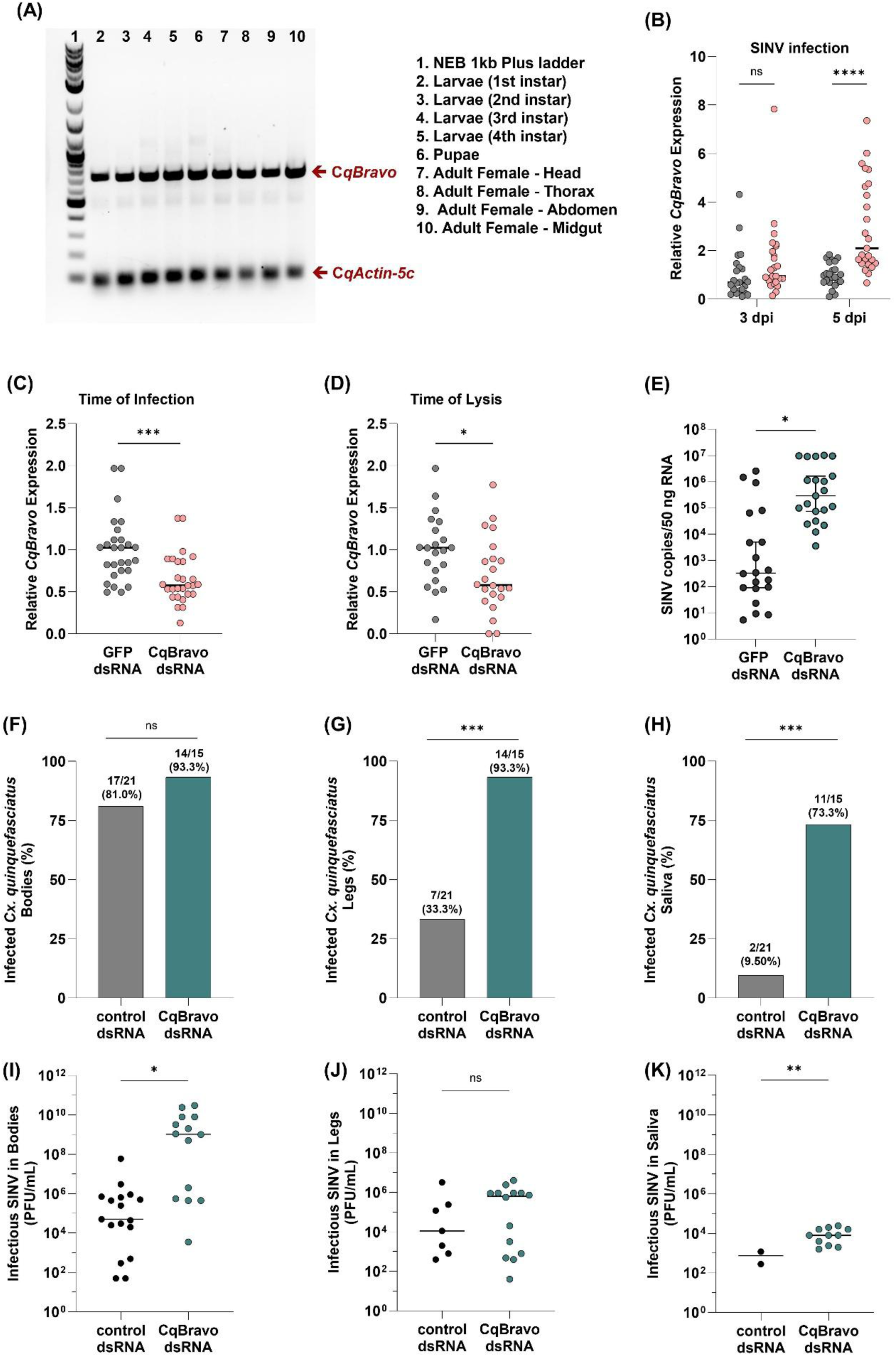
CqBravo is antiviral *in vivo* and upregulated following virus infection. *CqBravo* expression from *Cx. quinquefasciatus* tissues (5-7 day old females) and life stages was evaluated by RT-PCR (A). *CqBravo* expression was measured by RT-qPCR in whole mosquitoes following SINV infection (pink bars) at 3 dpi (n=23) and 5 dpi (n=25) compared to uninfected mosquitoes (grey bars) (B). Mosquitoes were injected intrathoracically with 1μg dsRNA targeting GFP (control) or CqBravo and silencing was confirmed by RT-qPCR at 3 days post injection (C; time of infection) and 8 days post injection (D; time of lysis). Injected mosquitoes were infected with SINV by infectious bloodmeal 3 days post injection, RNA was extracted at 5 dpi, and SINV RNA quantified by RT-qPCR (E). RNA quantification of *CqBravo* (B, C, D) was normalized to *CqActin-5c* and is presented relative to the GFP control. SINV RNA was quantified using a standard curve to determine copy numbers per 50ng of RNA (E). In a subsequent experiment, *Cx. quinquefasciatus* mosquitoes were injected with dsRNA as above, infected with SINV, and bodies (F, I), legs and wings (G, J), and saliva (H, K) were homogenized for plaque assays on Vero cells at 5 dpi. The percentage of positive SINV tissues is shown (F-H) and the SINV titers from tissues found to be positive for SINV (I-K). Significant changes in RNA abundance or virus titer compared to the control were analyzed by Welch’s t-test. Significant changes in the proportion of infected tissues (F-H) were analyzed by a Fishers Exact test. All P-values are shown as *P<0.05, **P<0.01, ***P<0.001, ****P<0.0001 (F-H).

To further investigate the antiviral role of CqBravo *in vivo,* we silenced *CqBravo* gene expression via intrathoracic injection of dsRNA targeting *CqBravo* mRNA or a control dsRNA targeting *GFP* and infected mosquitoes with SINV via infectious bloodmeal. At the time of infection, there was a significant (P <0.0001) 35% decrease in *CqBravo* gene expression (Figure 2C). At the time of lysis, there was still a significant (P<0.05) 30% decrease in *CqBravo* mRNA, but some expression had been restored (Figure 2D). When *CqBravo* expression was silenced, there was a significant 2,634-fold, increase in SINV copies (P<0.05), confirming that it is also antiviral *in vivo* (Figure 2E). To further define the antiviral impact of CqBravo on infectious virus titer and transmission potential, we next performed *CqBravo* silencing, SINV infection, and subsequent plaque assays from collected saliva, legs/wings, and mosquito bodies 5 dpi (Figure 2F-K). In the bodies of *CqBravo* silenced mosquitoes, 93.3% were positive for SINV infectious particles compared to 81.0% in control mosquitoes (Figure F). In *CqBravo* silenced mosquitoes, all infected mosquitoes (93.3%) were SINV positive in their legs and wings compared to only 33.3% of GFP control mosquitoes marking a significant (P <0.001) increase in dissemination of SINV in *CqBravo* silenced mosquitoes (Figure 2G). Similarly, 73.3% of saliva from *CqBravo* silenced mosquitoes was positive for SINV, while in control mosquitoes, only 9.50% of saliva sampes were SINV positive (P <0.001) (Figure 2H). When comparing SINV titers, *CqBravo* silencing resulted in a significant increase (P<0.05) in virus titers compared to the GFP control by 1,418 fold (Figure 2I). While in the legs and wings, there was no significant difference in SINV virus titers between the control and *CqBravo* silenced mosquitoes, there was a non-significant 1.6 fold trend (Figure 2J). In saliva of SINV infected mosquitoes there was a significant 13.5-fold increase in SINV titers in *CqBravo* silenced mosquitoes (Figure 2K). Overall, our results support that CqBravo is antiviral against SINV in *Cx. Quinquefasciatus* mosquitoes.

### Overexpression of CqBravo Reduces USUV Replication

Given that silencing *CqBravo* expression in Hsu cells reduced both LACV and USUV RNA levels, we wanted to determine the effects of *CqBravo* overexpression on arbovirus replication. Overexpression may provide stronger rationale for its future applicability in transmission control strategies. We thus transfected Hsu cells with a plasmid expressing FLAG-tagged *CqBravo* under the control of a polyubiquitin promoter and stained for FLAG to validate successful transfection, expression, and cellular localization of CqBravo (Figure 3A). *CqBravo* was readily expressed in transfected cells and localized primarily to the nucleus, despite the absence of a classic nuclear localization signal (NLS) in CqBravo (Figure 3A). We next transfected FLAG-CqBravo in parallel with an otherwise identical FLAG-GFP expressing control plasmid, and infected Hsu cells with either LACV or USUV (Figures 3B-E). We validated *CqBravo* overexpression using RT-qPCR and observed an 88-fold increase (P<0.0001) compared to the GFP control at the time of LACV infection (Figure 3B). We observed a small 23% decrease (P<0.05) in LACV RNA levels when *CqBravo* was overexpressed (Figure 3C). For USUV experiments, there was an 82-fold increase in *CqBravo* at the time of infection in FLAG-CqBravo transfected cells (Figure 3D), and *CqBravo* overexpression reduced USUV RNA by 73% (P <0.0001) compared to the control (Figure 3E).

**Figure 3.**
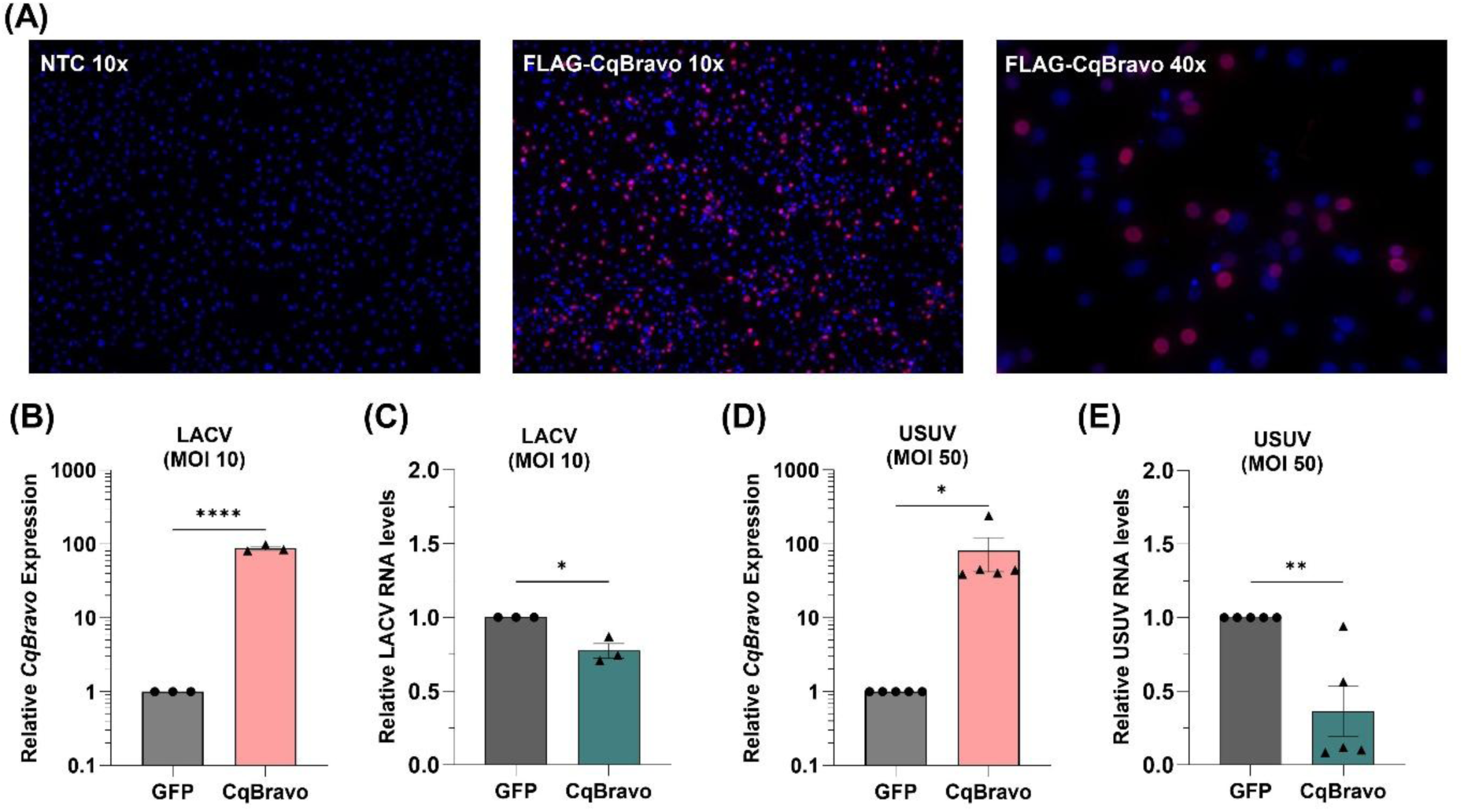
*CqBravo* overexpression reduces virus replication. Hsu cells were transfected with a plasmid expressing 3xFLAG-CqBravo under the control of a Pub promoter and expression was confirmed by immunofluorescence staining for FLAG (A). Cells were visualized using a Keyence BzX-710 microscope at 10x and 40x magnification as indicated (A). Hsu cells were transfected with either Pub-3xFLAG-GFP or Pub-3xFLAG-CqBravo, infected 48 hpt with either LACV at MOI 10 (B, C) or with USUV at MOI 50 (D, E), and RNA was extracted 2 dpi. *CqBravo* expression and viral RNA levels were measured by RT-qPCR, normalized to *CqActin5c*, and are shown relative to the GFP control. Each data point represents one biological replicate experiment with three technical replicates. Bars and error bars indicate the mean of three experiments with SEM. Significant changes in RNA abundance compared to the control were analyzed by one-tailed unpaired t-test (A-F). P-values are indicated as *P<0.05, **P<0.01, ***P<0.001, ****P<0.0001.

### CqBravo truncations reveal requirements for antiviral activity and nuclear localization

To gain insight into CqBravo’s antiviral activity, we next developed truncation constructs of CqBravo to identify which regions and domains (RRM1-3, PABP) in the protein may be critical for antiviral activity. Using our wild type (wt) CqBravo construct, we first made four different C-terminally truncated constructs that removed the unstructured C-terminus (Truncation#1, Figure 4A), as well as the RRM3 (Truncation#2, Figure 4A), the PABP domain (Truncation#3, Supplemental Figure 1A) or everything except the N-terminus and RRM1 (Truncation #4, Supplemental Figure 1A). We tested these constructs for their antiviral activity against USUV compared to the full-length CqBravo and found that only Truncation #1, with an intact RRM3, maintained antiviral activity that was not significantly different (P>0.05) from full-length CqBravo (Figure 4B, Supplemental Figure 1B), but there was a nonsignificant trend for mildly reduced activity of Truncation #1 (Figure 4B). Truncations #2 actually modestly increased USUV replication, possibly acting as a dominant negative construct (Figure 4B). We also performed N-terminal truncations that removed either only RRM1 (Truncation #5, Supplemental Figure 1A) or all of the N-terminal domains up to the start of RRM3 (Truncation #6, Figure 4A). Both of these constructs maintained antiviral activity that was not significantly different to full-length CqBravo with very similar levels of USUV reduction (Figure 4C, Supplemental Figure 1C). We also confirmed cellular expression and localization of our constructs and found that antiviral activity correlated with nuclear localization (Figure 4D, Supplemental Figure 1D).

**Figure 4.**
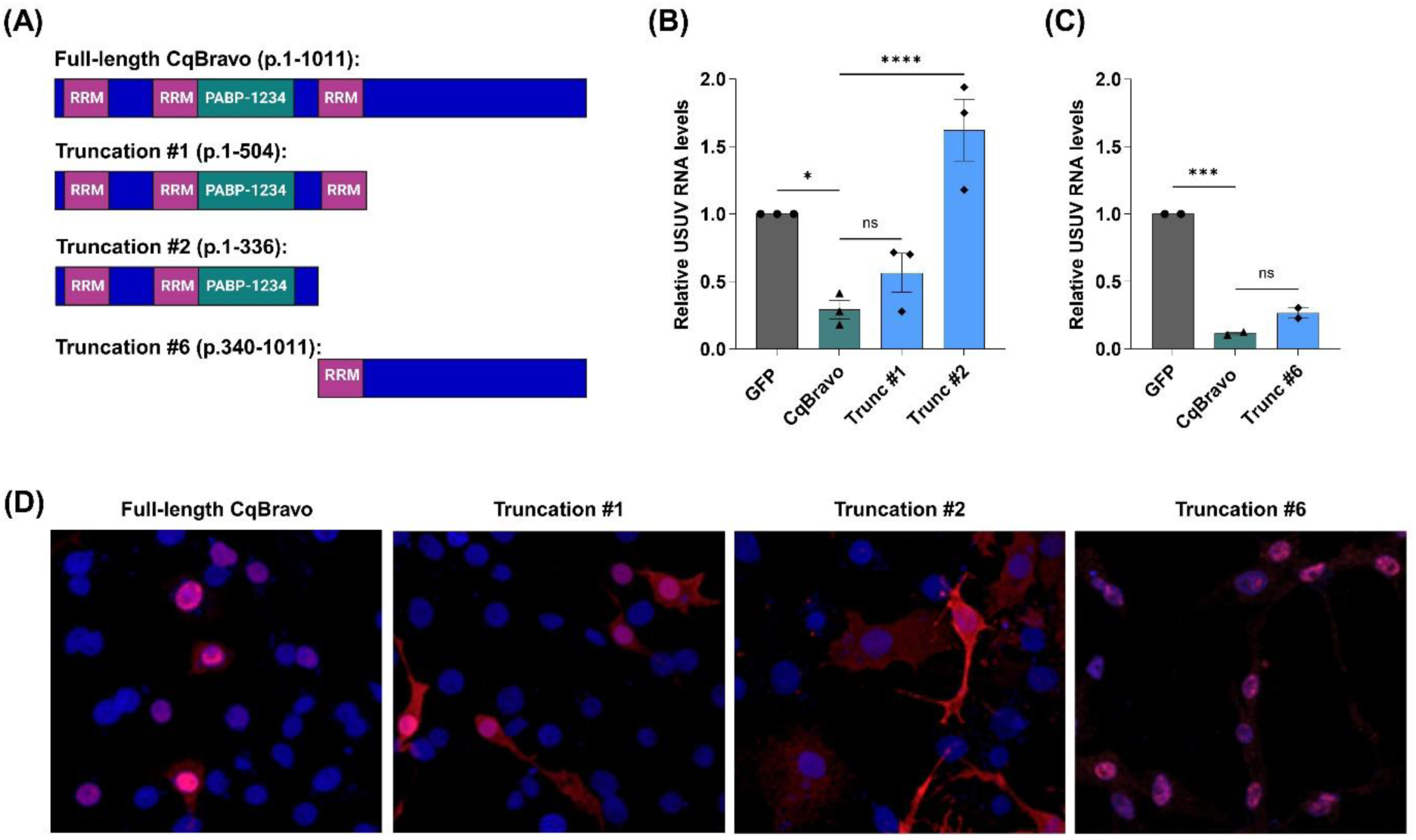
CqBravo determinants of antiviral activity and nuclear localization. CqBravo truncations were generated from the C-terminus and N-terminus of CqBravo (A) using the 3xFLAG-CqBravo expression plasmid. Hsu cells were transfected with GFP, full-length CqBravo, or truncations and infected 48 hpt with USUV at MOI 50. RNA was extracted 2 dpi, viral RNA was measured by RT-qPCR, normalized to *CqActin5c*, and are shown relative to the GFP control (B, C). In parallel, transfected and USUV infected Hsu cells were also stained using an anti-FLAG antibody to confirm CqBravo expression and localization (D). Cells were visualized using a Leica Stellaris 5 HyD S Confocal Microscope (D). Each data point (B, C) represents one biological replicate experiment with three technical replicates. Bars and error bars indicate the mean of three experiments with SEM. Significant changes in RNA abundance compared to full-length CqBravo were analyzed by one-way ANOVA (B, C). P-values are indicated as *P<0.05, **P<0.01, ***P<0.001, ****P<0.0001.

Truncation #1 retained partial nuclear localization with some cytosolic localization, while Truncations #2-4 localized almost exclusively to the cytosol (Figure 4D, Supplemental Figure 1D). Truncations #5 and #6 were almost exclusively localized to the nucleus (Figure 4D, Supplemental Figure 1D), again correlating with their antiviral activity (Figure 4C, Supplemental Figure 1C).

### CqBravo RRM3 Facilitates Antiviral Activity

Although CqBravo lacks a clear nuclear localization signal, RRM3 and/or the unstructured C-terminus of the protein seem important for nuclear localization. Given that RRM3 appears to be involved in facilitating nuclear localization and antiviral activity, we made another CqBravo truncation that consisted only of the 75 aa RRM3 domain (Figure 5A, Truncation #7). We also generated a CqBravo construct containing a mutated RRM3 domain (Figure 5B, RRM3_mut_). In RRMs, aromatic residues within the highly conserved ribonucleoprotein (RNP) domains, RNP1 and RNP2, were found to be critical to RNA binding activity (25, 26). We identified the RNP1 and RNP2 domains within RRM3 using AlphaFold structure prediction (27) and mutated all aromatic residues to alanines to remove RNA binding activity (Figure 4B). Since we do not know its RNA binding partners and there are two other RRMs, there was no simple way to validate the loss of RNA binding, but we expect loss of RRM3-RNA interactions based on the literature (25, 26). We transfected Hsu cells with constructs expressing Truncation #7 and CqBravo-RRM3_mut_, full-length CqBravo, and a FLAG-tagged GFP control, infected with USUV, and measured viral RNA 48 hpi (Figure 5C). CqBravo Truncation #7 decreased USUV RNA levels compared to the GFP control, but there was a significant increase compared to cells transfected with full-length CqBravo (P <0.001) resulting in an intermediate phenotype (Figure 5C). Expression of CqBravo-RRM3_mut_ also exhibited an intermediate phenotype with a decrease of USUV RNA compared to the GFP control, but a significantly reduced antiviral effect compared to full-length CqBravo (Figure 5C). These data show that the RRM3 domain significantly contributes to CqBravo antiviral activity, but RRM3 alone cannot maintain complete antiviral activity and loss of RRM3-RNA interactions may not result in a complete loss of antiviral activity. Both constructs were still able to localize to the nucleus (Figure 5C), possibly suggesting the presence of a cryptic NLS in RRM3.

**Figure 5.**
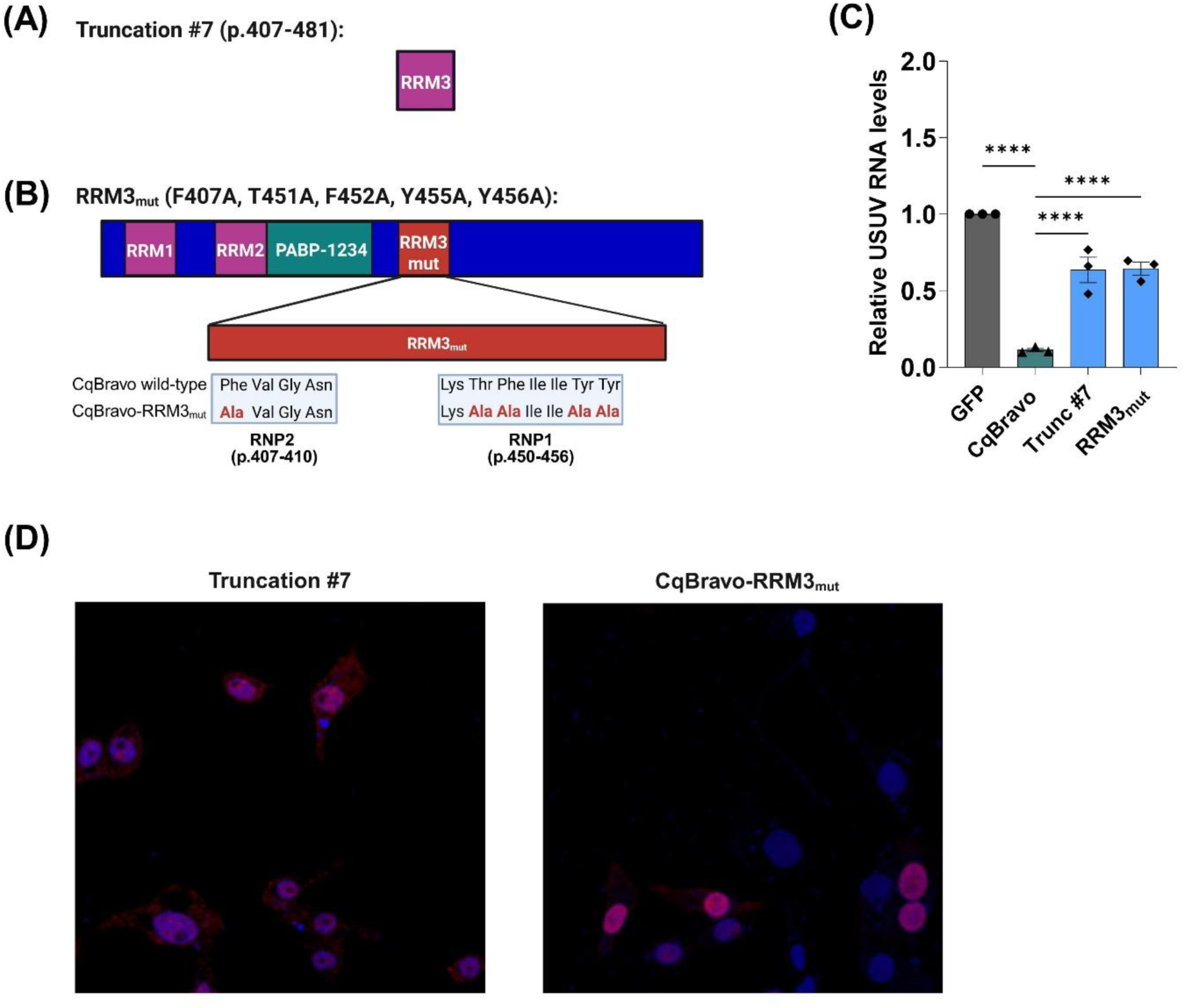
RRM3 is critical but not sufficient for CqBravo antiviral activity. A CqBravo truncation containing only the third RRM (RRM3) was generated from the 3xFLAG-CqBravo expression plasmid (A). In a full-length 3xFLAG-CqBravo construct, mutations were introduced in RRM3 of CqBravo, replacing key amino acids with aromatic ring side chains with alanines (B). Hsu cells were transfected with GFP, full-length CqBravo, the RRM3 truncation, or the RRM3 mutant (RRM3mut) and infected 48 hpt with USUV at MOI 50. RNA was extracted 2 dpi, viral RNA was measured by RT-qPCR, normalized to *CqActin5c*, and are shown relative to the GFP control (C). In parallel, transfected and USUV infected Hsu cells were also stained using an anti-FLAG antibody to confirm CqBravo expression and localization (D). Cells were visualized using a Leica Stellaris 5 HyD S Confocal Microscope (D). Each data point (C) represents one biological replicate experiment with three technical replicates. Bars and error bars indicate the mean of three experiments with SEM. Significant changes in RNA abundance compared to full-length CqBravo were analyzed by one-way ANOVA (B, C). P-values are indicated as *P<0.05, **P<0.01, ***P<0.001, ****P<0.0001.

### CqBravo overexpression alters the proteome of Hsu cells

Next, we sought to identify cellular pathways associated with CqBravo antiviral activity. We performed quantitative proteomic analysis of Hsu cells transfected with full-length CqBravo, Truncation #2, or a GFP control plasmid (n = 4/treatment group). Truncation #2 was selected because it lacks RRM3 and the C-terminal region of CqBravo and does not maintain antiviral activity against USUV (Figure 4). Across all samples, 5,118 proteins were detected and quantified (Supplemental Table 1). Principal component analysis (PCA) revealed that full-length CqBravo-expressing cells clustered separately from GFP- and Truncation #2-expressing cells, whereas GFP and Truncation #2 samples showed greater similarity and partial overlap (Figure 6A). As PCA demonstrated separation of treatment groups, we next used orthogonal partial least squares discriminant analysis (OPLS-DA) to identify proteins contributing to these differences (Figure 6B,C). The 30 proteins with the highest variable importance in projection (VIP) scores, representing the proteins that contributed most strongly to separation among treatment groups, are shown in Figure 6C together with their relative abundance across samples.

**Figure 6.**
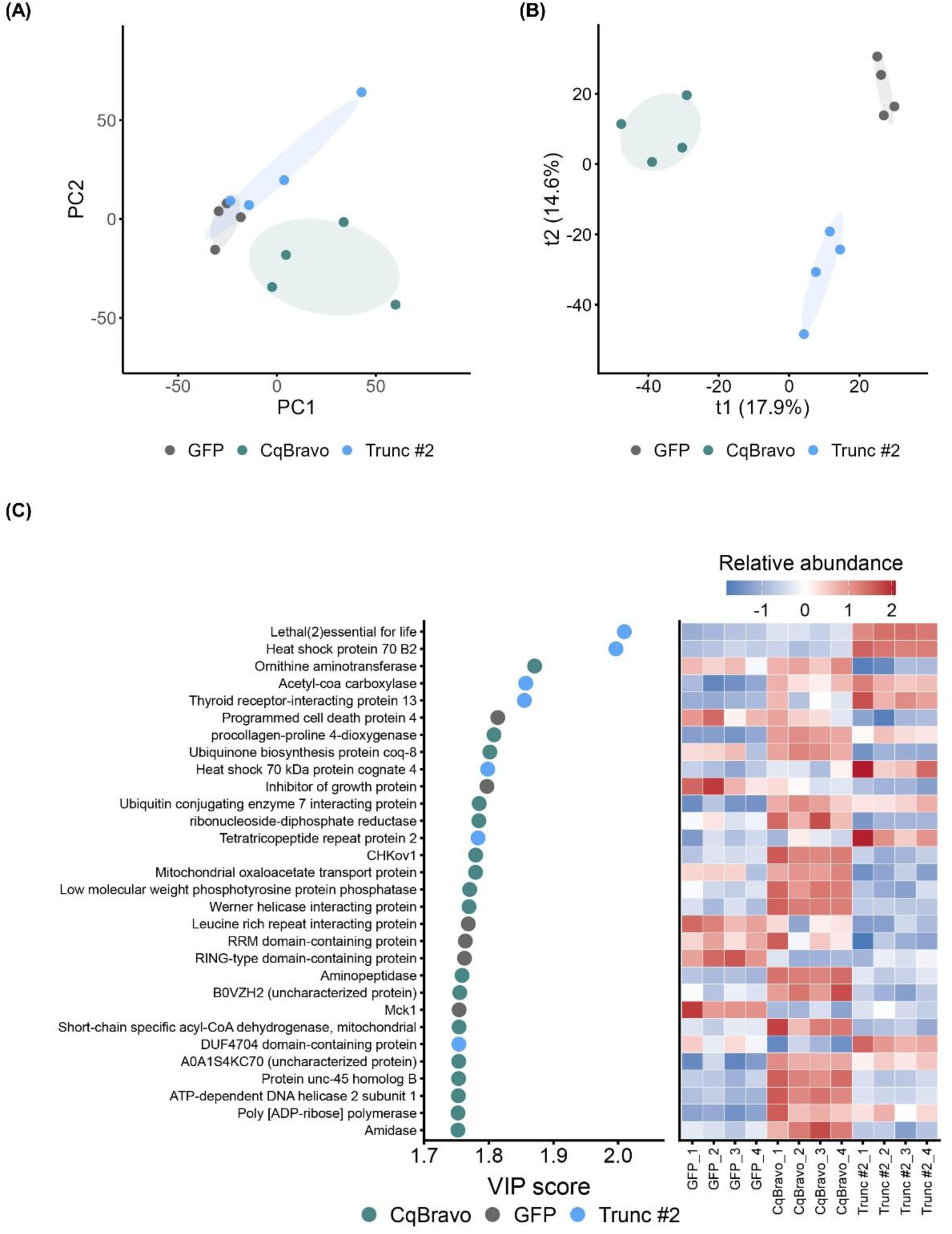
CqBravo expression alters the global proteomic landscape of Hsu cells. Hsu cells were transfected with full-length CqBravo, Truncation #2, or a GFP control plasmid. At 72 hpt, whole-cell lysates were collected, normalized to equivalent protein concentrations, and subjected to proteomic analysis. (A) Principal component analysis (PCA) of normalized protein abundance values. (B) Orthogonal partial least squares discriminant analysis (OPLS-DA) of proteomic profiles. (C) Variable importance in projection (VIP) scores for the top proteins contributing to separation of treatment groups in the OPLS-DA model, with the corresponding heatmap showing their relative abundance across samples.

Differential abundance analysis revealed 27 significantly altered proteins (24 upregulated, 3 downregulated) in cells expressing full-length CqBravo compared to GFP controls (Figure 7A). In contrast, only 2 proteins were significantly altered in Truncation #2-expressing cells relative to GFP controls (Figure 7B). Direct comparison of full-length CqBravo and Truncation #2 identified 14 significantly altered proteins (12 upregulated, 2 downregulated) in full-length CqBravo-expressing cells compared to Truncation #2-expressing cells (Supplemental Figure 2A). These represented a subset of the 27 proteins upregulated between CqBravo and GFP. When comparing abundance of all 29 significantly altered proteins across all three conditions, 16 proteins were only altered in full-length CqBravo overexpressing cells, but some proteins had an intermediate phenotype in Truncation #2 expressing cells (Supplemental Figure 2 B-DD). As expected, one of the proteins upregulated in both full-length CqBravo and Truncation #2 was CqBravo itself (Supplemental Figure 2H). Together, these findings demonstrate that full-length CqBravo and Truncation #2 are associated with distinct proteomic states and that removal of RRM3 and the C-terminal region substantially reduces the number of proteins that are differentially abundant relative to GFP controls.

**Figure 7.**
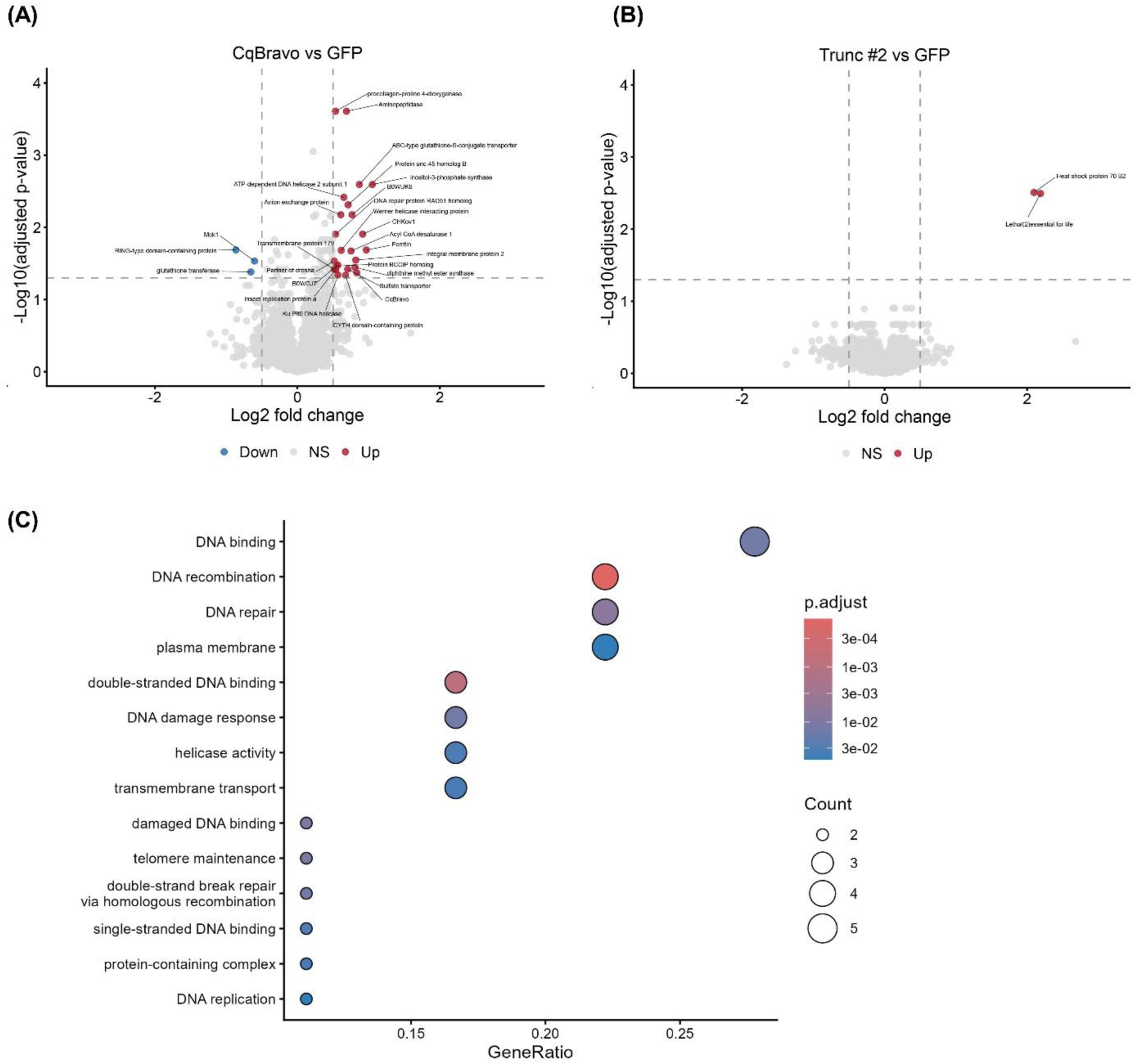
CqBravo expression is associated with altered abundance of proteins involved in DNA repair and genome maintenance. Hsu cells were transfected with full-length CqBravo, Truncation #2, or a GFP control plasmid. At 72 hpt, whole-cell lysates were collected, normalized to equivalent protein concentrations, and subjected to proteomic analysis. (A) Volcano plot showing differentially abundant proteins in full-length CqBravo-expressing cells relative to GFP controls. (B) Volcano plot showing differentially abundant proteins in Truncation #2-expressing cells relative to GFP controls. Proteins meeting significance thresholds (Benjamini-Hochberg adjusted P < 0.05 and |log2FC| ≥ 0.5) are highlighted. (C) Gene Ontology Biological Process enrichment analysis of proteins differentially abundant in full-length CqBravo-expressing cells relative to GFP controls. Significantly enriched terms (Benjamini-Hochberg adjusted P < 0.05) are shown. Dot size indicates the number of proteins associated with each term, and color indicates the adjusted P-value.

Finally, we sought to determine whether the proteomic changes observed between full-length CqBravo- and GFP-expressing cells were associated with specific cellular processes. Gene Ontology (GO) enrichment analysis identified significant enrichment of pathways involved in genome maintenance, including DNA repair, DNA metabolism, and ATP-dependent DNA processing (Figure 7C). Overall, we found that the regions of CqBravo required for antiviral activity are also associated with distinct changes in the host proteome, particularly in proteins involved in genome maintenance, shedding light on pathways that may contribute to CqBravo’s antiviral activity.

## Discussion

In this study, we investigated the antiviral role of the RNA-binding protein CqBravo in *Cx. quinquefasciatus* mosquitoes and derived Hsu cells. We found that CqBravo is highly expressed both in Hsu cells and at all life stages of *Cx. quinquefasciatus* development. We demonstrated that CqBravo is broadly antiviral, reducing replication of LACV and USUV in Hsu cells, and SINV in *Cx. quinquefasciatus* mosquitoes, and that it is upregulated during arbovirus infection in cell culture and *in vivo*. Additionally, we observed that CqBravo localizes to the nucleus in Hsu cells, despite the lack of a canonical NLS. Through the generation of CqBravo truncation and mutant constructs, we identified that its 3^rd^ RRM domain is important for nuclear localization and antiviral activity against USUV, but it may not be the sole component of CqBravo that facilitates antiviral activity. Finally, quantitative proteomic analysis revealed that CqBravo overexpression selectively altered the abundance of proteins involved in maintaining genome integrity, including DNA repair, recombination, replication, and helicase activity, providing the first mechanistic insight into the cellular pathways associated with CqBravo antiviral activity.

Bravo was initially identified in *Ae. aegypti* as a mosquito-specific protein that interacts with key components of the RNAi pathway, including Dicer-2, Ago2, and Piwi4, while acting independently of Dicer-2 activity (15). Our data supports that Bravo is also broadly antiviral in *Cx. quinquefasciatus*, and its function is conserved across mosquito species. However, we observed virus-specific differences in its antiviral effects, with CqBravo overexpression having a stronger effect on USUV replication than LACV replication.

*In vivo*, CqBravo silencing significantly increased SINV replication and dissemination in *Cx. quinquefasciatus* mosquitoes. The high percentage of virus dissemination to the legs, wings, and saliva in CqBravo silenced mosquitoes and the high viral titers observed in the bodies indicate a significant contribution to controlling virus replication, dissemination, and transmission *in vivo*. However, the impacts of CqBravo overexpression *in vivo* have yet to be tested using transgenic approaches and present an exciting new avenue for transmission control strategies.

On a cellular level, we show that exogenously expressed CqBravo localizes to the nucleus in *Cx. quinquefasciatus*-derived cells, despite lacking a canonical NLS. Through truncation analyses, we identified that its 3rd RRM domain (RRM3) is important for this nuclear localization. All truncations containing RRM3 localized predominantly to the nucleus, while all truncations lacking RRM3 localized mainly to the cytoplasm. RNA-binding proteins have been shown to contain non-classical nuclear localization signals (ncNLSs) that facilitate nuclear import (28, 29). While nuclear localization in other proteins is mediated through RRM binding within an RRM domain (30, 31), mutating RRM3 to disrupt RNA interactions still resulted in nuclear localization of CqBravo. It is possible that the mutations we introduced did not fully disrupt RRM3-RNA interactions, since we did not have any way to validate the disruption of RNA binding. However, based on previous studies, these mutations are predicted to disrupt residues critical for RNA binding and thereby impair RNA-binding activity (25, 26). Overall, it remains unclear whether CqBravo’s nuclear localization is directly mediated by an ncNLS, or other factors, such as interactions with host proteins or RNA itself.

Another important molecular finding was that CqBravo’s antiviral activity appears to rely on RRM3 and to a lesser extent, the unstructured C-terminus. None of the CqBravo truncations lacking RRM3 maintained antiviral activity. All truncations including RRM3, exhibited some level of antiviral activity, but only those constructs including the C-terminus of CqBravo exhibited antiviral activity that was nearly identical to full-length CqBravo. RRM3 alone was not sufficient to fully recapitulate the antiviral effects of the full-length protein, but given the short 75 aa peptide sequence, RRM3 expression could experience effects on folding or rapid degradation that prevent efficient antiviral activity. Similarly, mutation of the RNA-binding residues within RRM3 reduced Bravo’s antiviral activity but did not fully eliminate it, supporting the hypothesis that the RRM3 RNA-binding activity of CqBravo is critical but not solely responsible for antiviral activity. These data on the importance of RRM3 will be crucial as we further explore RNA-protein interactions, CqBravo’s cellular function, and the determinants of its antiviral activity.

Our proteomic analyses provide an additional perspective on potential mechanisms underlying CqBravo antiviral activity. Full-length CqBravo expression was associated with enrichment of proteins involved in genome maintenance, including DNA repair, DNA metabolism, and ATP-dependent DNA processing, whereas this enrichment was not present in cells transfected with the Truncation #2 plasmid which lacked the RRM3 domain and C-terminus of CqBravo. Although USUV and LACV replicate in the cytoplasm, both viruses encode proteins that interact with nuclear host processes. Specifically, arboviruses are known to manipulate DNA repair proteins, helicases, and other genome maintenance pathways to restrict host cellular processes and establish a cellular environment favorable for viral replication (32–38). For example, flavivirus capsid and NS5 proteins localize to the nucleus and are known to alter host transcriptional, antiviral, and DNA damage response pathways (40–42), while the orthobunyavirus NSs protein has been linked to host transcriptional shutoff and activation of DNA damage response pathways (46). Consequently, CqBravo’s antiviral activity may not arise through pathways that directly restrict viral replication. Instead, CqBravo may help preserve cellular pathways commonly targeted by arboviruses, creating a more refractory intracellular environment that is less permissive to productive infection.

The nuclear localization of CqBravo raises important questions about its mechanism of action. In *Ae. aegypti*, aBravo was shown to interact with RNAi pathway proteins, including Piwi4 and Ago2, but its direct role in RNAi-mediated antiviral defense remains unclear. In *Ae. aegypti* antiviral activity was independent of Dicer-2 activity, indicating aBravo functions outside of the canonical siRNA pathway in *Ae. aegypti* (15). Given its nuclear localization in *Cx. quinquefasciatus*, interactions with cytosolic proteins (e.g. Dicer-2, Ago-2, Piwi4) may not be critical for its activity, but may indicate indirect interactions through shared RNA binding or artificial interactions following cell lysis. It is also possible that CqBravo shuttles between the nucleus and cytosol, resulting in these interactions.

As a nuclear RNA-binding protein, CqBravo may participate in transcriptional regulation, mRNA transport, RNA stability, or RNA processing, including alternative splicing. Consistent with this possibility, our proteomic analyses identified enrichment of proteins involved in genome maintenance pathways, suggesting that CqBravo may influence these host nuclear processes. However, the direct molecular targets of CqBravo remain unknown. Therefore, identifying the RNAs bound by CqBravo will be critical for determining whether the genome maintenance pathways identified by proteomic analysis are direct targets of CqBravo-mediated regulation or arise indirectly as downstream consequences of CqBravo activity. Furthermore, future studies will be required to determine whether modulation of these pathways contributes to antiviral defense and whether manipulation of these pathways during CqBravo knockdown can recapitulate CqBravo’s antiviral phenotype.

Overall, our study provides evidence that CqBravo is an important antiviral factor in *Cx. quinquefasciatus* mosquitoes and mosquito-derived cells. Our findings suggest that CqBravo’s function is conserved across mosquito species and that its antiviral activity is broad across multiple arbovirus families. We also highlight the importance of the 3rd RRM domain in Bravo’s nuclear localization and antiviral activity setting the stage for future studies into its cellular role and mechanism of antiviral activity.

## Acknowledgements

The project described was supported by grants from the National Institute of Allergy & Infectious Diseases (R21AI185435) and Laura St. Clair was also supported by NIH/NIA individual LRP award L70AG084124.

## Notes

### Competing Interest Statement

The authors have declared no competing interest.

