## Supplemental Figures for "Bravo! A Mosquito Antiviral Protein that Restricts Arboviruses through an RRM3-Dependent Nuclear Mechanism"

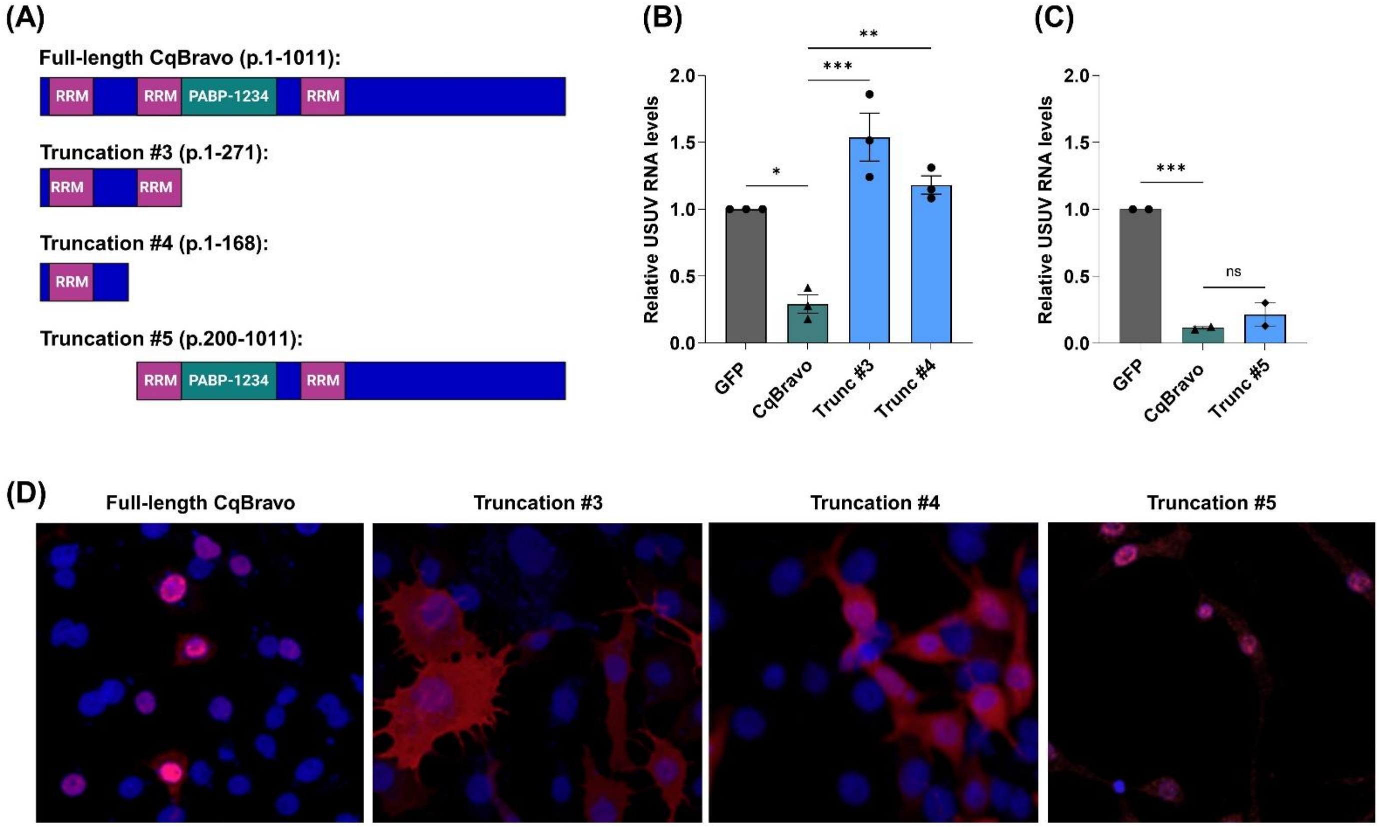


**Supplemental Figure 1. CqBravo truncations and their antiviral activity and cellular localization.** CqBravo truncations were generated from the C-terminus and N-terminus of CqBravo (A) using the 3xFLAG-CqBravo expression plasmid. Hsu cells were transfected with GFP, full-length CqBravo, or truncations and infected 48 hpt with USUV at MOI 50. RNA was extracted 2 dpi, viral RNA was measured by RT-qPCR, normalized to *CqActin5c*, and are shown relative to the GFP control (B, C). In parallel, transfected and USUV infected Hsu cells were also stained using an anti-FLAG antibody to confirm CqBravo expression and localization (D). Cells were visualized using a Leica Stellaris 5 HyD S Confocal Microscope (D). Each data point (B, C) represents one biological replicate experiment with three technical replicates. Bars and error bars indicate the mean of three experiments with SEM. Significant changes in RNA abundance compared to full-length CqBravo were analyzed by one-way ANOVA (B, C). P-values are indicated as *P<0.05, **P<0.01, ***P<0.001, ****P<0.0001.

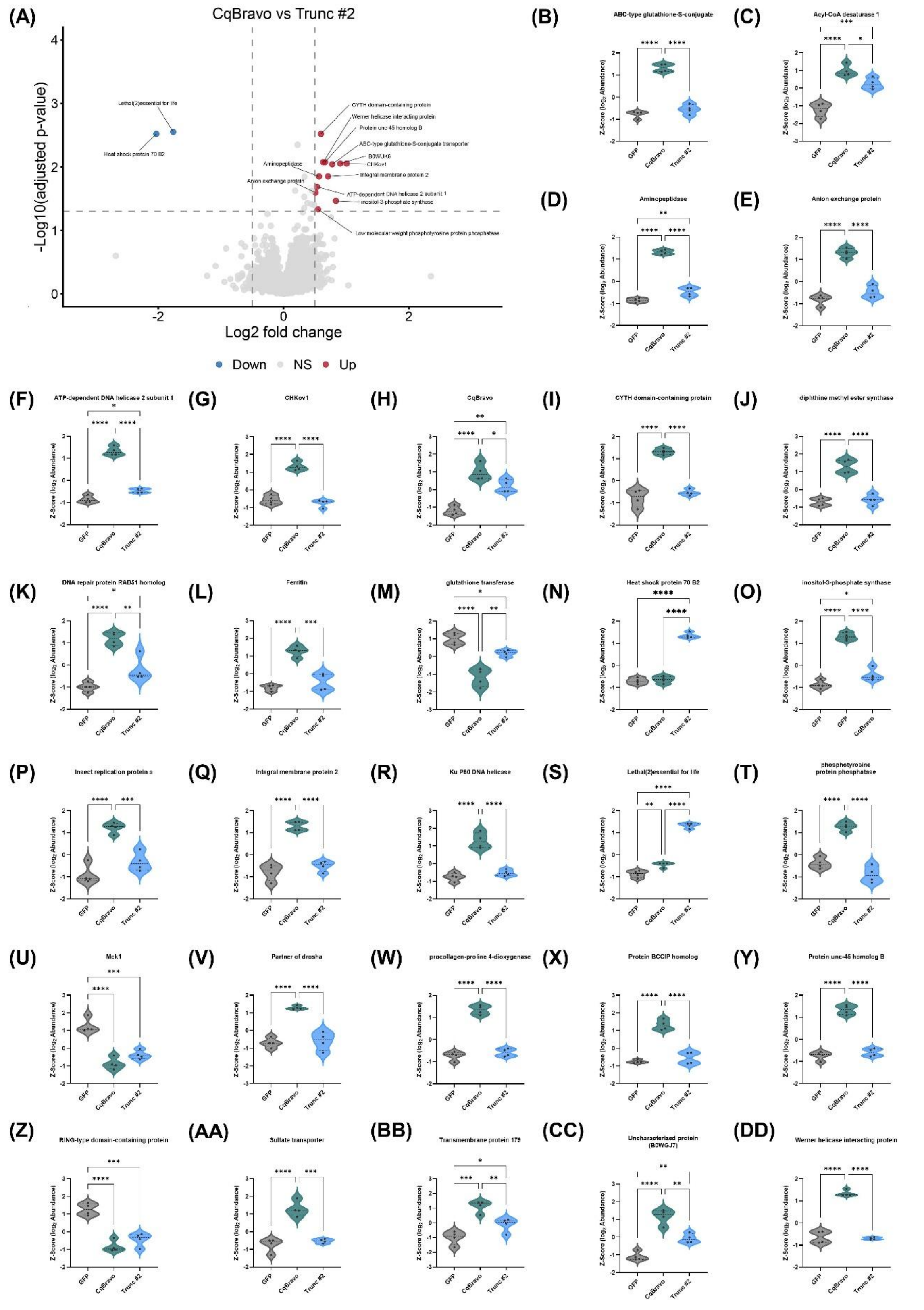


**Supplemental Figure 2. Relative abundance of proteins identified as differentially abundant among GFP, wt CqBravo, and Truncation #2-expressing HSU cells.** Hsu cells were transfected with wt CqBravo, Truncation #2, or a GFP control plasmid. At 72 hpt, whole-cell lysates were collected, normalized to equivalent protein concentrations, and subjected to proteomic analysis. (A) Volcano plot showing differentially abundant proteins in wt CqBravo-expressing cells relative to Truncation 2-expressing cells. Proteins meeting significance thresholds (Benjamini-Hochberg adjusted $P < 0.05$ and $|\log2FC| \geq 0.5$) are highlighted. (B-DD) Violin plots showing relative abundance of proteins identified as significantly differentially abundant between full-length CqBravo and GFP or Truncation #2 and GFP. Protein abundance values were log2-transformed and Z-score normalized within each protein. Statistical significance was assessed by one-way ANOVA with Tukey's multiple-comparisons testing. Significance is indicated as *$P < 0.05$, **$P < 0.01$, ***$P < 0.001$, and ****$P < 0.0001$.
